# A systematic approach for identifying shared mechanisms in epilepsy and its comorbidities

**DOI:** 10.1101/269860

**Authors:** Charles Tapley Hoyt, Daniel Domingo-Fernández, Nora Balzer, Anka Güldenpfennig, Martin Hofmann-Apitius

**Author notes:** **Corresponding Author:** Hoyt, C. T. Department of Bioinformatics, Fraunhofer Institute for Algorithms and Scientific Computing, Sankt Augustin 53754, Germany. Telephone details. These authors contributed equally to this work (Charles Tapley Hoyt, Daniel Domingo-Fernández).

## Abstract

Cross-sectional epidemiological studies have shown that the incidence of several nervous system diseases is more frequent in epilepsy patients than in the general population. Some comorbidities (e.g., Alzheimer’s disease and Parkinson’s disease) are also risk factors for the development of seizures; suggesting they may share pathophysiological mechanisms with epilepsy.

A literature-based approach was used to identify gene overlap between epilepsy and its comorbidities as a proxy for a shared genetic basis for disease, or genetic pleiotropy, as a first effort to identify shared mechanisms. While the results identified neurological disorders as the group of diseases with the highest gene overlap, this analysis was insufficient for identifying putative common mechanisms shared across epilepsy and its comorbidities. This motivated the use of a dedicated literature mining and knowledge assembly approach in which a cause-and-effect model of epilepsy was captured with Biological Expression Language.

After enriching the knowledge assembly with information surrounding epilepsy, its risk factors, its comorbidities, and antiepileptic drugs, a novel comparative mechanism enrichment approach was used to propose several downstream effectors (including the GABA receptor, GABAergic pathways, etc.) that could explain the therapeutic effects carbamazepine in both the contexts of epilepsy and AD.

We have made the Epilepsy Knowledge Assembly available at https://www.scai.fraunhofer.de/content/dam/scai/de/downloads/bioinformatik/epilepsy.bel and queryable through NeuroMMSig at http://neurommsig.scai.fraunhofer.de. The source code used for analysis and tutorials for reproduction are available on GitHub at https://github.com/cthoyt/epicom.

## Introduction

Seizures are transient occurrences of signs and symptoms due to abnormal or excessive neuronal activity in the brain [1]. Classically, their underlying causes were thought to be the primary drivers for increasing mortality in epilepsy. While epilepsy has been classically studied as a disorder of the brain characterized by an enduring predisposition to epileptic seizures, it is no longer considered a condition in which seizures are the only concern [2]. Epilepsy is also associated with several comorbidities, including Alzheimer’s disease (AD), Parkinson’s disease (PD), other nervous system diseases, and psychiatric disorders [3-5], due to a variety of genetic, biological, and environmental factors [6].

The prevalence of migraine in epilepsy patients (under 64 years old) is 5.71% in contrast to 3.47% in the general population [7]. The mechanistic understanding of epilepsy and migraine presumes that they share the underlying pathophysiology related to alterations in sodium and calcium ion channels and ion transporters (sodium-potassium transport) [8,9]. Moreover, drugs acting on voltage-gated sodium channels and γ-aminobutyric acid (GABA) receptors (e.g., valproate, topiramate, etc.) are used not only prevent migraine attacks, but are also used as antiepileptic drugs [10,11].

The prevalence of epileptic seizures in AD patients is strongly influenced by genetic factors [12] — epilepsy and seizures occur more often in patients with early-onset familial AD than those with sporadic AD [13]. Further, convulsive seizures have been described in approximately 40% of familial AD patients with the PSEN1 p.Glu280Ala mutation [14], 30% with PSEN2 mutations [15], and 57% with amyloid precursor protein (APP p.Thr174Ile) duplications [16]. Additionally, the p.Pro86Lys mutation in CALHM1 (rs11191692) is associated with both AD and temporal lobe epilepsy through its influence on calcium homeostasis [17].

Evidence that some comorbidities (e.g., AD and depression) also act as risk factors for developing seizures suggests they may share pathophysiological mechanisms [13, 18]. Conversely, epileptic seizures (as well as neoplasms) have been reported to cause intellectual disabilities in patients with tuberous sclerosis (Jansen *et al*., 2008; Osborne *et al*., 2010). Furthermore, certain antiepileptic drugs (e.g., topiramate) are associated with higher incidence of cognitive problems [1, 21].

The several causal and associative relationships observed between epilepsy and other indications on the phenotypic level warrant further investigation for shared elements on the genetic and molecular levels. While inquiry on the genetic level often begins with genome-wide association studies to identify shared loci, identifying the appropriate data set(s) and linking intergenic single nucleotide polymorphisms (SNP) to their functional consequences across scales in complex disease is still a significant challenge [22]. Even after identifying disease-associated genes, it is difficult to assess their individual contributions to the complex etiology of epilepsy and its related indications. Thus, capturing the different causal relationships between biological entities involved in the pathophysiology of a complex disease is an essential step to a better understanding of the processes that lead to the disease state.

Here, we present two methods to hypothesize shared mechanisms: first, a literature-based approach for quantifying gene overlap between epilepsy and its comorbidities as a proxy for a shared genetic basis of disease, or genetic pleiotropy; and second, a systematic approach using the NeuroMMSig mechanism enrichment server. Finally, we use these methods to propose an explanation for the observed therapeutic effects of a drug that has been studied in the contexts of epilepsy and AD. A schematic representation of the methodology, analysis, and results is presented in **Figure 1**.

**Figure 1:**
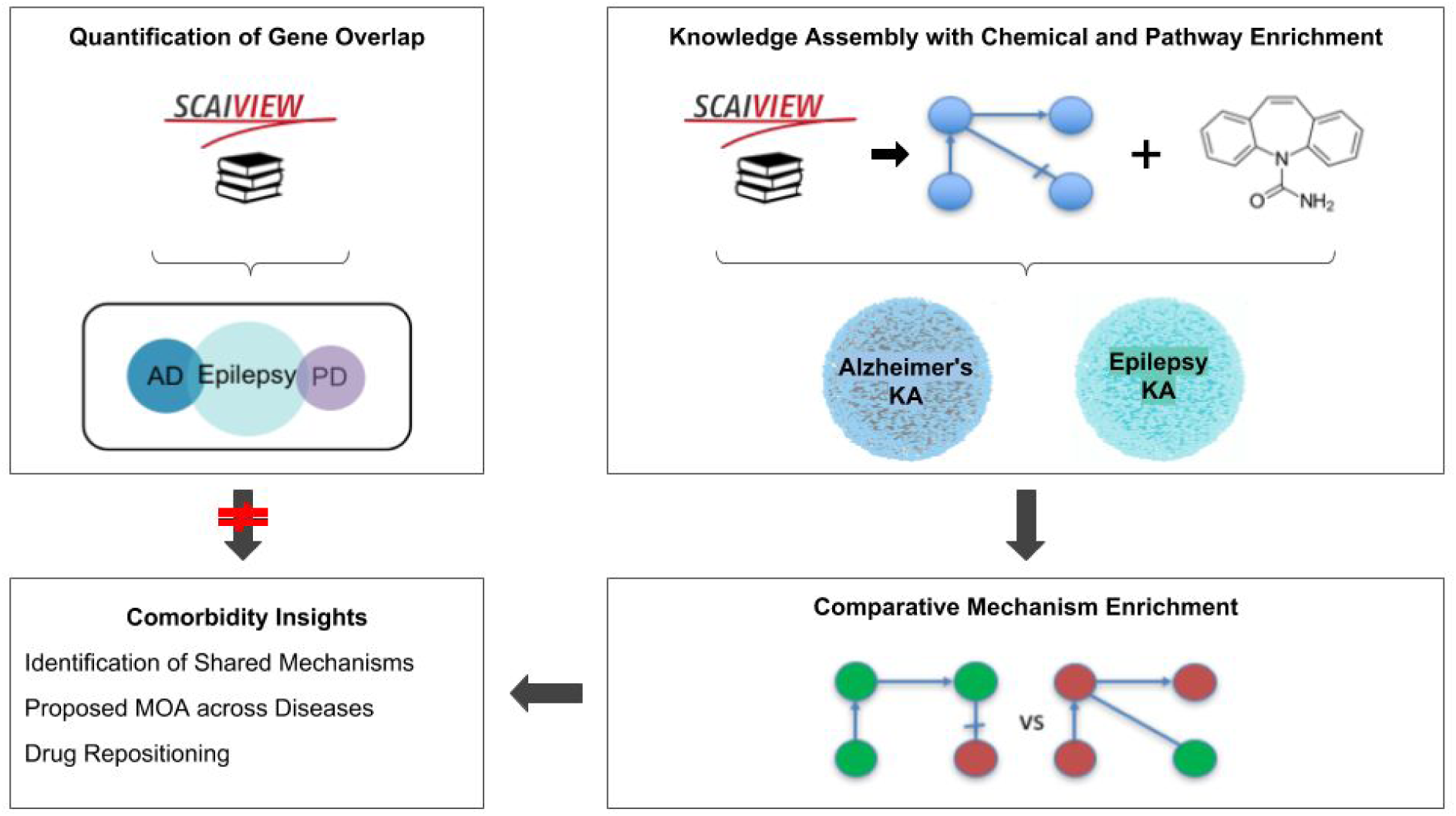
A graphical abstract of the methodology, analysis, and results presented in this work. The two upper boxes represent the methodology while the two lower boxes represent the analysis and results. The upper-left box outlines the quantification of gene overlap between epilepsy and its well-known comorbidities described in Keezer *et al.* [23] (e.g., Alzheimer’s disease, Parkinson’s disease, etc.) using literature based methods. The upper-right box outlines the assembly of knowledge from epilepsy literature with chemical and pathway enrichment as described in the “Preparation for Mechanism Enrichment” and “Relation Extraction” sections. The lower-right box represents the comparative mechanism enrichment that was used to generate comorbidity insights (lower-left box) after literature-based methods proved insufficient.

## Materials and Methods

### Quantification of Gene Overlap

SCAIView v1.7.3, indexed from MEDLINE on 2016-07-14, (http://academia.scaiview.com/academia) was used to identify and quantify the overlap of genes co-occurring in the literature with epilepsy and its novel and well-known comorbidities reviewed by Keezer *et al.* [23]. At least one disease was selected for each of the ICD (International Statistical Classification of Diseases and Related Health Problems) groups that were analyzed by Keezer *et al.* [23] as well as the addition of Parkinson’s disease due to its previously published relevance [24].

In order to later assess publication bias in literature-based methods, we report the total number of documents associated with each disease by querying SCAIView with their corresponding MeSH terms and the total number of associated genes that have a positive relative entropy (i.e., occur more frequently in the results of a given SCAIView query than in the rest of the SCAIView indexed literature) in the context of the query as described by Younesi *et al*. [25] Gene sets for each comorbidity were then retrieved by constructing queries using their corresponding MeSH terms joined with the epilepsy MeSH term by the “AND” operator. The associated genes for each comorbidity query were identified with the same method and the epilepsy pleiotropy rate was calculated as the percentage of the genes associated with the comorbidity query in the set of associated genes with epilepsy **(Table 1)**.

**Table 1:** Results of the Epilepsy comorbidity analysis using SCAIView. The number of ***associated documents*** (column 2) retrieved from SCAIView for each disease are shown given a reference query using corresponding the MeSH term from column 1. The ***disease-associated genes*** (column 3) contains the number of genes relevant to the corpus retrieved from a disease-specific query. The ***comorbidity-associated genes*** (column 4) contains the number of genes relevant to the comorbidity query between the target disease and epilepsy. Lastly, the ***epilepsy pleiotropy rate*** (column 5) describes the ratio of the count of genes reported in column 4 with the total number of epilepsy-associated genes (2901).

| Disease (MeSH ID) | Associated documents | Disease associated Genes | Comorbidity associated genes | Epilepsy pleiotropy rate (%) |
| --- | --- | --- | --- | --- |
| Epilepsy (D004827) | 192245 | 2901 | - | - |
| Alzheimer's Disease (D000544) | 109495 | 4968 | 396 | 13.65 |
| Migraine (D008881) | 30928 | 1230 | 306 | 10.54 |
| Parkinson's Disease (D010300) | 79103 | 3646 | 258 | 8.89 |
| Hypertension (D006973) | 391190 | 5574 | 252 | 8.68 |
| Dementia (D003704) | 183802 | 5833 | 220 | 7.58 |
| Diabetes Mellitus (D003920) | 394411 | 6661 | 184 | 6.34 |
| Anxiety (D001007) | 84138 | 1782 | 124 | 4.27 |
| Arthritis (D001168) | 259327 | 5367 | 122 | 4.20 |
| Cataract (D002386) | 52150 | 2238 | 119 | 4.10 |
| Colonic Neoplasms (D003110) | 107274 | 3646 | 30 | 1.03 |
| Urinary Incontinence (D014549) | 34170 | 720 | 24 | 0.82 |
| Peptic Ulcer (D010437) | 68234 | 1445 | 21 | 0.72 |
| Pulmonary Disease, Chronic Obstructive (D029424) | 35627 | 2244 | 15 | 0.51 |

For example, 226 genes were found to be associated with diabetes using the comorbidity query `[MeSH Disease:”Epilepsy”] AND [MeSH Disease:”Diabetes Mellitus”]`. Of these, 184 had positive relative entropies, which indicate that these genes occur more frequently in literature mentioning both diseases than in the rest of the SCAIView indexed literature. Finally, the epilepsy pleiotropy rate was calculated normalizing 184 by the total number of genes (2901) with a positive relative entropy found by querying for epilepsy `[MeSH Disease:”Epilepsy”]`, resulting in a pleiotropy rate of 6.34%.

### Relation Extraction

While literature co-occurrence can generate initial hypotheses about genetic pleiotropy, it does not provide sufficient mechanistic insight to explain the clinical observations in epilepsy and its comorbidities. The increasing quantity of knowledge in the biomedical domain makes it difficult or impossible for researchers to be knowledgeable in any but incredibly specific topics [26]. The task of manually generating pleiotropy hypotheses that explain overlap between the aetiological mechanisms of epilepsy and its comorbidities is daunting. In order to enable computer-aided automatic reasoning, the knowledge surrounding epilepsy and its comorbidities was systematically extracted from the literature using manual relation extraction and encoded in Biological Expression Language (BEL) [27].

First, a corpus was generated from the 192245 documents related to epilepsy retrieved (**Table 1**) to further investigate the causal relations surrounding the identified genes. A second corpus was generated from the 2666 documents retrieved by querying SCAIView for “epilepsy” and its sub-terms in the Epilepsy Ontology [28] occuring with the free text, “comorbidity. “Manual relation extraction and encoding in BEL was then performed starting with a select subset of the two corpora based on their prioritization by SCAIView to generate the Epilepsy Knowledge Assembly. The knowledge assembly was further enriched with pharmacological knowledge surrounding 19 antiepileptic drugs and their targets from the Pharmacogenetics and Pharmacogenomics Knowledge Base (PharmGKB) [29]. Ultimately, the knowledge assembly comprised relations from 641 unique citations. Finally, the PyBEL framework [30] was used to parse and validate the syntax and semantics of the underlying BEL Script. A summary of the contents of the Epilepsy Knowledge Assembly is presented in **Table 2**.

**Table 2:**
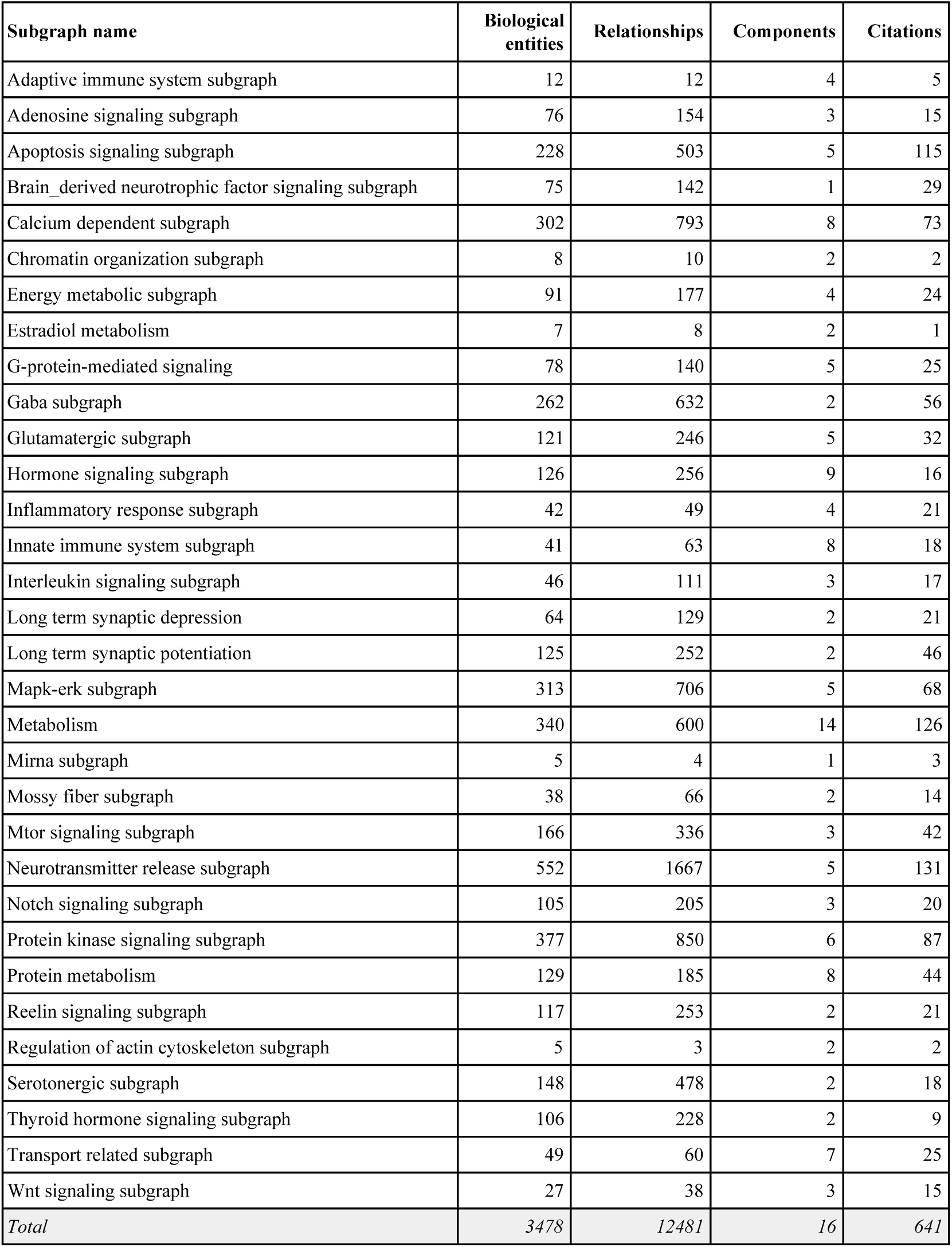
Statistics over the Epilepsy Knowledge Assembly, grouped by subgraph generated by PyBEL v0.10.0. The biological entities entry corresponds to the number of genes, chemicals, proteins, biological process, etc. in a subgraph. The number of relationships (i.e., edges) corresponds to the number of publications in which a relationship was described. The number of components corresponds to the number of “connected” graphs exist within a subgraph. Finally, the number of citations corresponds to the total number of articles that were curated in the subgraph. A more detailed summary is included in the supplementary information.

| Subgraph name | Biological entities | Relationships | Components | Citations |
| --- | --- | --- | --- | --- |
| Adaptive immune system subgraph | 12 | 12 | 4 | 5 |
| Adenosine signaling subgraph | 76 | 154 | 3 | 15 |
| Apoptosis signaling subgraph | 228 | 503 | 5 | 115 |
| Brain_derived neurotrophic factor signaling subgraph | 75 | 142 | 1 | 29 |
| Calcium dependent subgraph | 302 | 793 | 8 | 73 |
| Chromatin organization subgraph | 8 | 10 | 2 | 2 |
| Energy metabolic subgraph | 91 | 177 | 4 | 24 |
| Estradiol metabolism | 7 | 8 | 2 | 1 |
| G-protein-mediated signaling | 78 | 140 | 5 | 25 |
| Gaba subgraph | 262 | 632 | 2 | 56 |
| Glutamatergic subgraph | 121 | 246 | 5 | 32 |
| Hormone signaling subgraph | 126 | 256 | 9 | 16 |
| Inflammatory response subgraph | 42 | 49 | 4 | 21 |
| Innate immune system subgraph | 41 | 63 | 8 | 18 |
| Interleukin signaling subgraph | 46 | 111 | 3 | 17 |
| Long term synaptic depression | 64 | 129 | 2 | 21 |
| Long term synaptic potentiation | 125 | 252 | 2 | 46 |
| Mapk-erk subgraph | 313 | 706 | 5 | 68 |
| Metabolism | 340 | 600 | 14 | 126 |
| Mirna subgraph | 5 | 4 | 1 | 3 |
| Mossy fiber subgraph | 38 | 66 | 2 | 14 |
| Mtor signaling subgraph | 166 | 336 | 3 | 42 |
| Neurotransmitter release subgraph | 552 | 1667 | 5 | 131 |
| Notch signaling subgraph | 105 | 205 | 3 | 20 |
| Protein kinase signaling subgraph | 377 | 850 | 6 | 87 |
| Protein metabolism | 129 | 185 | 8 | 44 |
| Reelin signaling subgraph | 117 | 253 | 2 | 21 |
| Regulation of actin cytoskeleton subgraph | 5 | 3 | 2 | 2 |
| Serotonergic subgraph | 148 | 478 | 2 | 18 |
| Thyroid hormone signaling subgraph | 106 | 228 | 2 | 9 |
| Transport related subgraph | 49 | 60 | 7 | 25 |
| Wnt signaling subgraph | 27 | 38 | 3 | 15 |
| *Total* | *3478* | *12481* | *16* | *641* |

### Preparation for Mechanism Enrichment

The Epilepsy Knowledge Assembly was enriched with mechanistic annotations following the procedures outlined by Domingo-Fernández *et al.* [24] in order to integrate it into NeuroMMSig and enable multi-modal mechanism enrichment analyses with queries over genes, SNPs, and neuroimaging features.

A taxonomy of epilepsy mechanisms was generated by combining the list of well-established epilepsy mechanisms from Staley [31] with concepts from the Pathway Terminology System [32] co-occurring in articles matching either [MeSH Disease:”Epilepsy”] or entries in the Epilepsy Ontology indexed by SCAIView. The resulting 784 terms representing mechanisms were curated in order to normalize entities, remove irrelevant entries, and group similar terms. Next, relations in the Epilepsy Knowledge Assembly were annotated with mechanisms based on whether their entities were involved in the mechanism as outlined by Domingo-Fernández *et al.* [24]. During the annotation process, new mechanisms not yet included in the inventory were found in the literature; thus, the mechanism inventory was parallely updated until concluding the annotation with a total of 32 annotated subgraphs **(Table 2)**. More details and examples about the mapping procedure can be found on the NeuroMMSig introduction page.

In the next section, we use NeuroMMSig to identify shared mechanisms between epilepsy and AD on the basis of the mechanism of action of multi-indication drugs.

## Results and Discussions

### Investigation of Comorbidities

Of the 2901 genes identified as associated with epilepsy by having a positive relative entropy score, **Table 1** shows that nervous system disorders had the highest gene overlap (10.54% co-occurred with migraine, 7.58% with dementia, 8.89% with PD, and 13.65% with AD). These rates do not positively correlate with previous epidemiological studies of comorbidity incidence with the same conditions reviewed by Keezer *et al.* [23], therefore, literature-based gene overlap is a poor proxy for comorbidity incidence. These findings are plotted in supplementary **Figure S2**. For example, epilepsy patients under the age of 64 years showed the highest incidence of migraine and over the age of 64 years had the highest incidence of dementia [33] — both of which are inconsistent with the trend observed in literature co-occurrence. Interestingly, neither of these trends correlated with the number of disease-specific literature, which reflects publication bias.

While gene-centric methods alone were insufficient for unraveling the shared pathophysiology in epilepsy and studied comorbidities, systematic approaches for identifying and evaluating shared mechanisms may explain the aggregate effects of their interactions that cannot be captured by simple measures like literature co-occurrence.

### Mechanism Enrichment

Alzheimer’s disease was chosen as a putative comorbidity of epilepsy for further investigation not only because it had the highest literature-based gene overlap with epilepsy (**Table 1**), but additionally because of the prior existence of the Alzheimer’s Disease Knowledge Assembly [34] and its inclusion in NeuroMMSig.

Of the most frequently mentioned drugs in epilepsy literature **(Figure S1)**, we identified carbamazepine as having both notable target representation in the Alzheimer’s Disease Knowledge Assembly along with multiple references indicating its positive effects on memory and treatment of elderly patients with seizures [35] as well as positive effects in the treatment of AD patients [35,36].

In the following sections, we first use NeuroMMSig to investigate the mechanisms enriched by the targets of carbamazepine in the context of epilepsy. After, we make a comparative enrichment to identify possible overlapping mechanisms with AD in order to explore the therapeutic effects of the drug in both disease contexts.

#### Epilepsy Mechanism Enrichment

While carbamazepine has been observed to act through inhibition of sodium and calcium voltage-gated channels as well as activation of the GABA receptors [37, 38], its mechanism of action is still not fully understood [39]. In order to better understand the impact of the drug, the gene set of all of its known targets (**Text S1**), was queried on the NeuroMMSig mechanism enrichment server against the Epilepsy Knowledge Assembly.

Because the NeuroMMSig mechanism enrichment algorithm only provides relative scores, the top 10th percentile of results were used. Of the twelve networks with at least one mapped gene, those with an enrichment score in the top 10th percentile (above 0.696) were the adenosine signaling subgraph and the GABA subgraph (**Table S1**). While the increase of the purine nucleoside, adenosine, has been associated with the incidence of seizures, its mechanistic connection is still unknown [40]. However, recent research has identified several promising targets that regulate and balance adenosine levels such as adenosine kinase (ADK) and its receptors (ADORA family) [40, 41]. Similarly to adenosine, the inhibitory neurotransmitter response induced by GABA is responsible for balancing many excitatory signals occurring in the brain. Studies that investigated reduction and abnormalities in GABA-inhibitory processes lead to the development of GABA agonists (e.g., vigabatrin and tiagabine) that act as anticonvulsants in epilepsy patients [42].

After assessing the plausibility of these subgraphs’ involvement in the aetiology of epilepsy, the union of the networks was used to further investigate the relation between the downstream effects of carbamazepine and its therapeutic effect on epilepsy patients. Due to the its size, it was necessary to filter and query the network by finding the shortest paths between the drug’s targets and different biological processes of interest in the context of epilepsy then combining them to form a new graph to support quickly identifying candidate pathways. Finally, the common upstream controllers between each pathway were included to provide further context to their overlap. Combining, reasoning over, and manually interpreting the generated paths lead to the simplified network depicted in **Figure 2** representing the downstream effects of carbamazepine.

**Figure 2:**
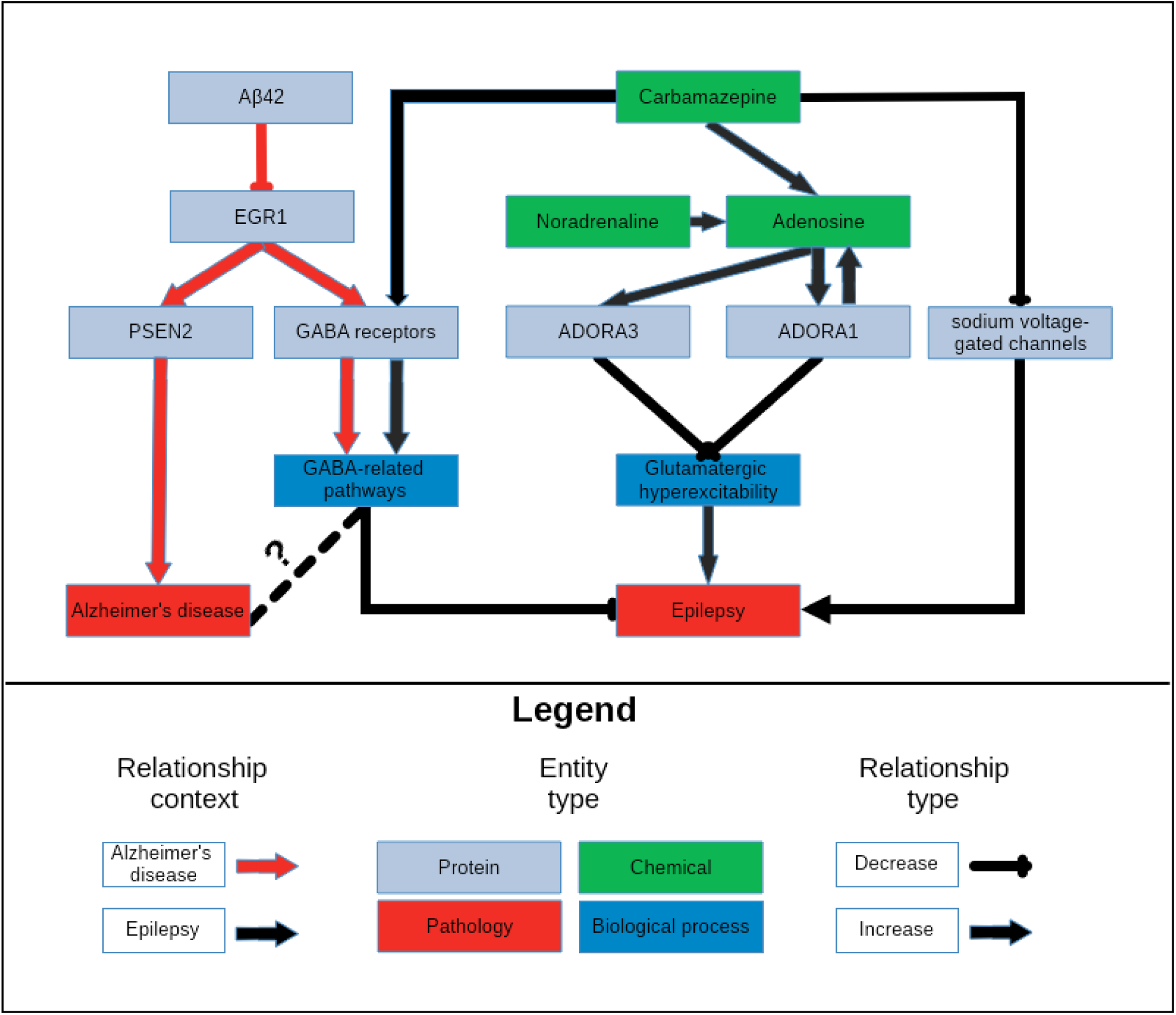
A schematic representation of the knowledge surrounding carbamazepine retrieved by querying its targets with NeuroMMSig. The relevant portions of the most significantly enriched graphs in the context of epilepsy and AD, the adenosine signalling and the GABA subgraphs, were merged and displayed in order to highlight a potential explanatory mechanisms for the therapeutic effects of carbamazepine (**Text S2)**. It is rendered with a hierarchical layout to mirror the flow from molecular entities to proteins, biological processes, and pathologies.

The figure demonstrates the synergistic effects of the activation of the GABA receptor family and the inhibition of the sodium voltage-gated channels to potentiate GABA-mediated inhibition, and therefore, decrease the risk of developing seizures. Furthermore, carbamazepine causes an increase in the production of adenosine, whose receptors, ADORA1 and ADORA3, in a positive feedback loop with the production of adenosine which ultimately leads to a reduction in glutamatergic excitatory signals. Synaptic plasticity in response to the downstream effects of these signals may contribute to drug resistance, and ultimately, seizure recurrence.

#### Comparative Mechanism Enrichment

NeuroMMSig was queried with the same gene set in the context of AD in order to perform a comparative investigation of shared mechanistic perturbations with epilepsy. Because the mechanism of action of carbamazepine is poorly represented in the literature (**Table S2**) and its known targets are less implicated in AD, it was unsurprising that fewer subgraphs were enriched in the context of AD.

The most significant, the GABA subgraph, which describes the upstream controllers of the GABA receptor and its downstream effectors, highly overlapped with the GABA subgraph in the context of the epilepsy – it contains key relations that may explain the efficacy of carbamazepine in both conditions. Studies in AD models have shown a negative correlation between the abundance of amyloid beta 42 (Aβ42) and the expression of the transcription factor for the GABA receptor family EGR1 [43,44]. This correlation could be caused by an unknown controller in the aetiology of AD; in which state, a patient would have decreased expression of EGR1, and therefore fewer GABA receptors and less ability to inhibit the excitatory signals that lead to seizures. While the link between epilepsy and GABAergic neurotransmission has already been exploited by antiepileptic drugs, its role in AD is not yet clearly understood **(Figure 2)**. However, several recent publications have rationalized targeting GABAergic neurotransmission for treatment of AD [45, 46].

Tangentially, EGR1 upregulates the expression of PSEN2, a member of the catalytic subunit of the γ-secretase complex that regulates APP cleavage [47]. Mutations in PSEN2 that have been both linked to amyloid beta accumulation [48] and seizures in AD patients [49] provide further evidence for the existence for a shared mechanism through which carbamazepine acts in the contexts of AD and epilepsy **(Figure 2)**.

Noradrenaline is a known anticonvulsant [50] that is often lacking in patients in the early stages of AD due to an observed loss of noradrenergic neurons [51]. Because carbamazepine has been observed to activate noradrenergic neurons [52], its anticonvulsant activity may be due to it indirectly increasing noradrenaline levels. Finally, noradrenaline potentiates the previously mentioned adenosine pathways [53].

While mechanism enrichment provides several insights, a cursory search of PubMed of “Carbamazepine”[nm] AND “Alzheimer disease”[MeSH Terms] suggested publications [54,55] that implicate autophagy in the therapeutic action of Carbamazepine. Further investigation showed Carbamazepine does not have a significant effect on expression of proteins in the mTOR-pathway [54] and that it is likely increasing autophagic flux through an mTOR-independent pathway [55]. Mechanism enrichment analysis was unable to prioritize autophagy pathways due to some of the shortcomings of knowledge-based methods. For example, the AD NeuroMMSig subgraph corresponding to autophagy pathways did not contain any of the targets of Carbamazepine listed by PharmGKB. This could be due to the choice of boundaries in subgraph definition, or also due to the lack of annotation of autophagy-related targets in PharmGKB. Autophagy has been implicated in epilepsy [56], but the literature has not yet succinctly described the connection from the therapeutic to its target, pathway, and finally the pathology. Finally, because knowledge assemblies are inherently incomplete, this shows the complementary nature of the two approaches.

While the exact mechanism of action of carbamazepine remains elusive, this proposed mechanism enrichment approach was able to identify multiple pathways through which it could be acting in both the AD, epilepsy, and shared context. Looking forward, this approach can be applied across a wide variety of chemical matter in the neurodegenerative disease space, as well as in other domains for which appropriately annotated knowledge assemblies exist, in order to support identification of drugs’ mechanisms of actions, drug repositioning opportunities, and the development of new lead compounds.

## Conclusion

Our findings indicate that literature-based methods as a proxy for genetic pleiotropy generally do not correlate with the results from epidemiological studies of epilepsy and its comorbidities. Furthermore, strictly gene-centric methods lack the ability to elucidate mechanistic insight that a knowledge assembly can support.

After formalizing a representative sample of the knowledge surrounding epilepsy, its risk factors, its comorbidities, and antiepileptic drugs, we annotated mechanistic subgraphs to include in the NeuroMMSig mechanism enrichment server. Finally, an enrichment approach focusing on the targets of carbamazepine proposed the several downstream effectors (including the GABA receptor, GABAergic pathways, etc.) that could explain its therapeutic effects in both the contexts of epilepsy and AD.

Future work will include applying this procedure to a wider variety of drugs and chemical matter across different diseases. Finally, we have made the Epilepsy Knowledge Assembly publicly available through NeuroMMSig (http://neurommsig.scai.fraunhofer.de) to facilitate further systems biology and chemoinformatics investigations of epilepsy.

## Authors’ Contributions

C.T.H. and D.D.F. conceived and designed the study, implemented and interpreted the mechanism enrichment approach in NeuroMMSig, and wrote this manuscript. N.B. and A.G curated and annotated the Epilepsy Knowledge Assembly. M.H.A. reviewed the content.

## Acknowledgements

The authors would like to thank our colleagues at Fraunhofer SCAI for their critical review of this manuscript.

