## Supplementary Materials for "A systematic approach for identifying shared mechanisms in epilepsy and its comorbidities"

### Supplement

#### Supplementary Tables

| Subgraph name | Enrichment score |
|---|---|
| Adenosine signaling subgraph | 0.75 |
| GABA subgraph | 0.72 |
| Glutamatergic subgraph | 0.48 |
| Neurotransmitter release subgraph | 0.46 |
| Serotonergic subgraph | 0.33 |
| Notch signaling subgraph | 0.20 |
| Mossy Fiber Subgraph | 0.14 |
| Brain_derived neurotrophic factor signaling subgraph | 0.08 |
| Hormone signaling subgraph | 0.05 |
| Long term synaptic potentiation | 0.03 |
| Calcium dependent subgraph | 0.02 |
| Protein kinase signaling subgraph | 0.01 |

**Table S1.** Mechanism enrichment scores obtained by querying NeuroMMSig with the targets of carbamazepine in the context of epilepsy. Only subgraphs with an enrichment score in the top 10th percentile (above 0.696) were selected for further analysis.

| Search engine | Query | Articles retrieved |
|---|---|---|
| SCAIView | ([MeSH Disease:"Alzheimer Disease"]) AND [NDD:"GABAergic"] | 232 |
| SCAIView | ([MeSH Disease:"Epilepsy"]) AND [NDD:"GABAergic"] | 2043 |
| PubMed | (GABA receptor) AND Alzheimer | 266 |
| PubMed | (GABA receptor) AND Epilepsy | 3902 |

**Table S2: A comparison of GABA receptor literature in epilepsy and Alzheimer's disease literature.** SCAIView and PubMed were used to query the occurrence of GABA-related publications in the contexts of AD and epilepsy. The results show a much larger representation in epilepsy literature than in AD literature.

### Supplementary Figures

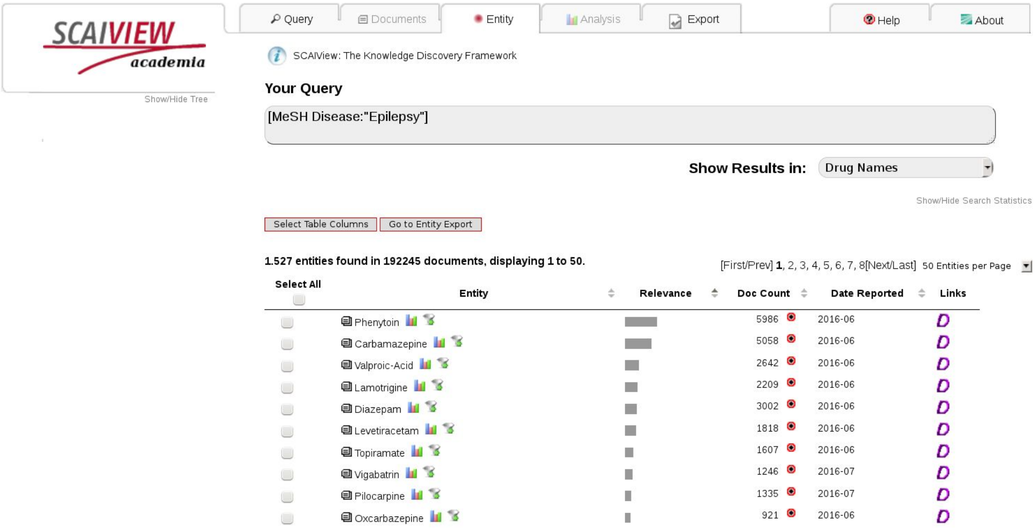


**Figure S1:** The most frequently mentioned drugs in epilepsy literature.

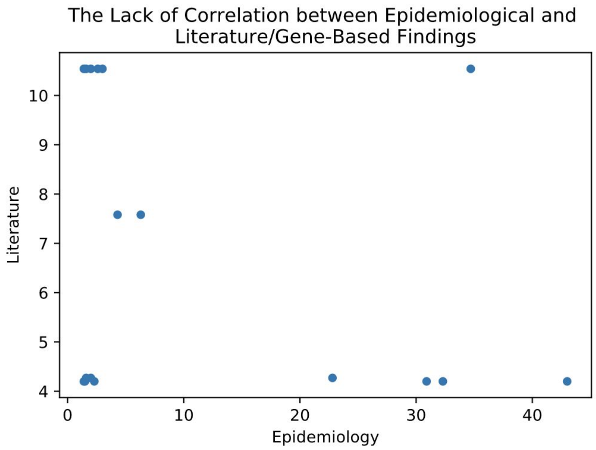


**Figure S2**: The epidemiological comorbidity incidences and literature-based gene overlaps have no correlation, so literature-based gene overlap is not a good proxy for comorbidity. Image generated by a Jupyter Notebook at https://github.com/cthoyt/EpiCom/blob/master/Epidemiology%20versus%20Literature-Based%20Comorbidity.ipynb.

### Supplementary Text

**Text S1: Genes used to query NeuroMMSig and their enriched subgraphs**

The gene set composed of the targets of carbamazepine consists of SCN1A, GABRA1, GABRA2, GABRA3, GABRA4, GABRA5, GABRA6, GABRB1, GABRB2, GABRB3, GABRG1, GABRG2, GABRG3, GABRD, GABRE, GABRP, GABRQ, ADORA1, and ADORA3.

**Text S2: List of articles (PubMed IDs) from which evidences to create Figure 2 were taken**

Epilepsy pathophysiology (black edges): 3280560, 16895979, 17054941, 18093662, 20831750, 22379998, 23535492, and 2553432.

Alzheimer's disease pathophysiology (red edges): 14585504, 19909279, 21969301, and 24919190.

**Text S3: Manual Crafting of the Mechanistic Subgraphs**

A detailed description of the mapping procedure between relationships and epilepsy related mechanisms can be found in the introduction page of NeuroMMSig.

**Text S4: Epilepsy Knowledge Assembly**

The Epilepsy Knowledge Assembly can be accessed under the following URL:

https://www.scai.fraunhofer.de/content/dam/scai/de/downloads/bioinformatik/epilepsy.bel
